# Microphysiological Modeling of Vascular Adipose Tissue for Multi-Throughput Applications

**DOI:** 10.1101/2024.01.30.578061

**Authors:** Michael Struss, Evangelia Bellas

## Abstract

Adipose tissue (AT) is a highly vascularized endocrine organ which regulates whole-body metabolic homeostasis. Key AT functions which rely on vascularization include insulin-stimulated glucose uptake and lipolysis (lipid mobilization to supply energy). Most in vitro AT models do not include the vasculature, and while microphysiological systems (MPS) incorporate spatial organization of cells, 3D environments, and perfusion by external pumps, they are too large to fit traditional cultureware. Thus, we developed a novel miniaturized vascularized adipose tissue (µAT) platform compatible with traditional 24 well plates. Using this µAT platform, we quantified vascular permeability and adipocyte function by insulin-stimulated glucose uptake and lipolysis assays. Shear flow decreased vascular permeability and increased insulin-stimulated glucose uptake. Treatment with forskolin, an adenyl cyclase agonist, increased lipolysis, and decreased vascular permeability. This µAT platform allows for the facile screening of compounds in a physiologically relevant system where both adipocyte and vascular function can be evaluated.

## Introduction

Adipose tissue (AT), previously thought of as a lipid storage unit, is now recognized as a highly vascularized endocrine organ, secreting biologically active signals into neighboring vasculature for crosstalk with other tissues[1]. This crosstalk supports key adipose tissue functions, such as insulin-stimulated glucose uptake and lipolysis, regulating glucose and lipid metabolism, neuroendocrine function, reproduction, and cardiovascular function[2],[3]. AT endocrine function requires close interactions with the vasculature, with almost every adipocyte in contact with a neighboring blood vessel[4]. In obesity, metabolic dysfunction primarily occurs as the result of adipocyte hypertrophy, without concomitant increases in vascularization such that AT becomes hypoxic, leading to pro-fibrotic and pro-inflammatory signaling[5]. Interactions between adipocytes and vasculature are needed for maintaining the proper AT function; therefore, modeling this crosstalk is crucial to study metabolic (dys)function.

The widespread adoption of increased throughput analytical methods has improved the efficiency and pace of biological research. While accelerating experimental workflows, these methods often rely on two dimensional (2D) or three dimensional (3D) cultures, which fail to mimic several aspects of the spatiotemporal organization and complexity of organ systems[6]. Conceptually, organs are composed of hundreds of “mini tissues” that respond to soluble factors and mechanical cues such as compression, shear flow, extracellular matrix (ECM) microarchitecture, binding motifs, and markers unique to specific cell populations[7]. Yet, many in vitro AT models do not include vasculature nor a perfusable vascular conduit to mimic this critical feature required for metabolic homeostasis and crosstalk. Appropriate 3D and environmental cues are important in developing AT models[8–18].

Microphysiological systems (MPS), or “organs-on-chips,” are platforms which more accurately model tissue microenvironment through spatially organized cell cultures, 3D microenvironments, and perfusable channels to apply stimuli, such as shear flow, to model high-level tissue complexity. MPS platforms have been developed to study various biological phenomena such as angiogenesis, tumor metastasis, and immune cell migration[6,19–23]. Yet many MPS platforms have not been adapted to traditional multi-throughput workflows as they are often too large for well plates or require external pumps or motors, limiting their multi-throughput potential.

To overcome these limitations, we have developed a novel miniaturized adipose tissue (µAT) platform, compatible with the traditional flat bottom 24 well plates, to model the interactions between adipocytes and their vasculature. This includes vascular function, such as vascular permeability and adipocyte function, by insulin-stimulated glucose uptake and lipolysis within µAT platform by sampling from the vessel compartment directly. Using miniaturized tissue compartments, subtractive molding fabrication techniques, and oscillatory gravity-driven flow, the platform is easily integrated into multi throughout well plates providing a low-cost, low resource tissue platform for easy integration with various multi-throughput applications.

## Results

### Morphological characterization of µAT platforms

First, we characterized the organization of adipocytes and vessel within the µAT platforms. The homogeneous organization of adipocytes after 7 days within the bulk hydrogel region in µAT platforms was confirmed via confocal microscopy (figure 2(b)). The µAT vessel within the bulk hydrogel was clearly visible with strong CD31 signal (figure 2(b)). Maximum intensity z projections of different layers (figure 2(c) – 2(e)) and orthogonal views (figure 2(f)) of the confocal image z-stacks confirmed an open lumen structure containing a confluent monolayer of endothelial cells.

**Figure 1.**
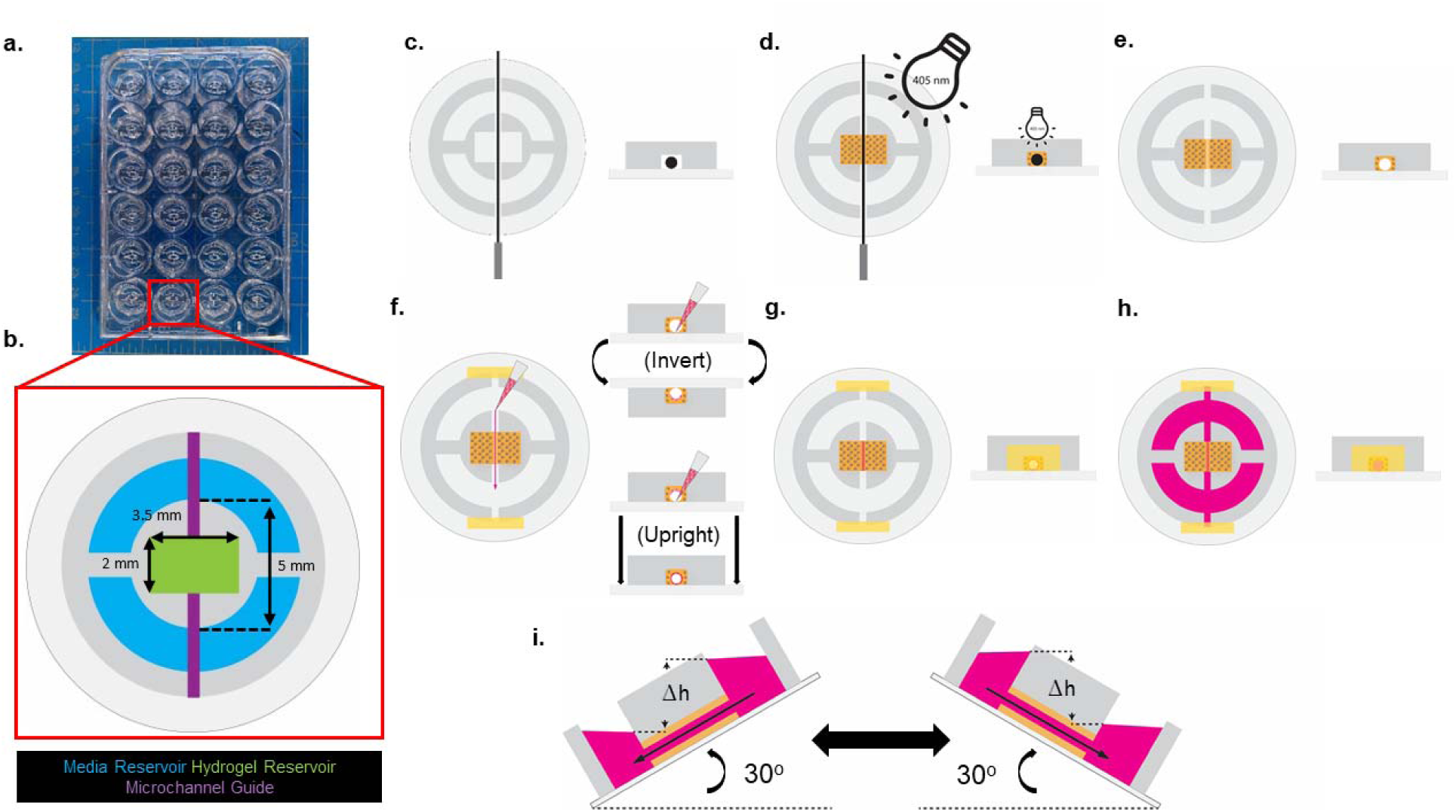
µAT Platform Fabrication. (**a)** µAT platforms were designed to be compatible with standard flat bottom 24 well plates and feature **(b)** two media reservoirs (blue), one hydrogel reservoir (green), and one microchannel guide (purple). Two media reservoirs allow for flow across the microchannel. The hydrogel reservoir contains the adipocytes encapsulated in a gelatin methacryloyl (GelMA) hydrogel. The microchannel guide aids in the fabrication process of the endothelial cell vessel. **(c)** A stainless-steel needle inserted through the µAT platform via the microchannel guide. **(d)** The µAT platform filled with a gelatin methacryloyl hydrogel solution containing adipocytes and crosslinked for 2 minutes using 405 nm light. **(e)** The stainless-steel rod is removed from the µAT platform, leaving a hollowed microchannel void, and the media reservoirs are sealed from outside to prevent leakages. **(f)** Endothelial cell suspension is added to each media reservoir, allowing for cells to flow through and adhere to the microchannel for 1 hr. This process is repeated 2X times to allow endothelial cells attachment to the top and bottom of the channel. **(g,h)** Endothelial cell suspension aspirated, washed 3X with 1X PBS, filled with fresh culture media and cultured in static or flow conditions. **(i)** Flow conditions were achieved by culturing the µAT platform on a biorocker with a 30° maximum rocking angle.

**Figure 2.**
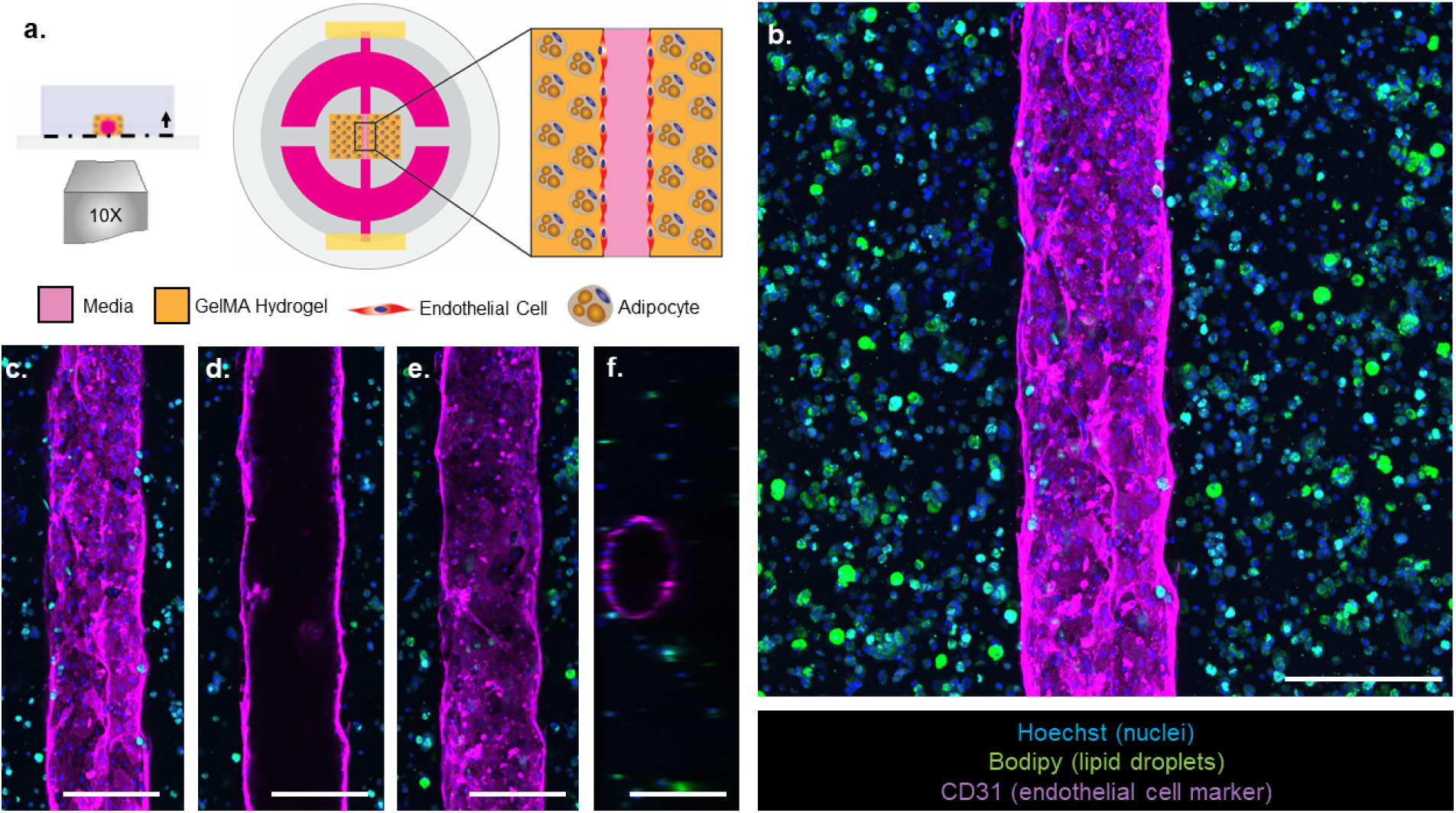
**Morphological Characterization of µAT Platforms**. **(a)** Experimental workflow for obtaining confocal z-stacks at the center of the µAT platform. **(b)** Maximum z-projection of the µAT platform with adipocytes stained with BODIPY 493/503 (green, lipid droplets), endothelial cells stained with CD31 (magenta, endothelial cell marker), and cell nuclei stained with Hoechst 33342 (blue, nuclei). **(c-f)** Individual slices from the z-stack show the **(c)** bottom, **(d)** middle, **(e)** top, and **(f)** orthogonal view of the endothelialized vessel within the µAT platform demonstrating the vessel is open lumen structure. Scale bars, 400 µm.

### Vessel permeability within µAT platforms

Vessel permeability of the vessel within µAT platforms was measured by perfusing media containing fluorescent dextran through the vessel via a hydrostatic pressure differential (figure 3(a)) and measuring the intensity in the vessel *(Io)* and hydrogel (∂*I/*∂*t)* regions to calculate the permeability coefficient (*Pd*) (figure 3(b) – 3(d)). On day 3, vascular permeability of µAT platforms were measured for all conditions. In static conditions, there were no significant differences in vessel permeability in the presence of endothelial cells compared to the adipocyte-only group (figure 3(e)). In the presence of shear flow, vessel permeability significantly decreased in the group with endothelial cells when compared to the adipocyte-only group.

**Figure 3.**
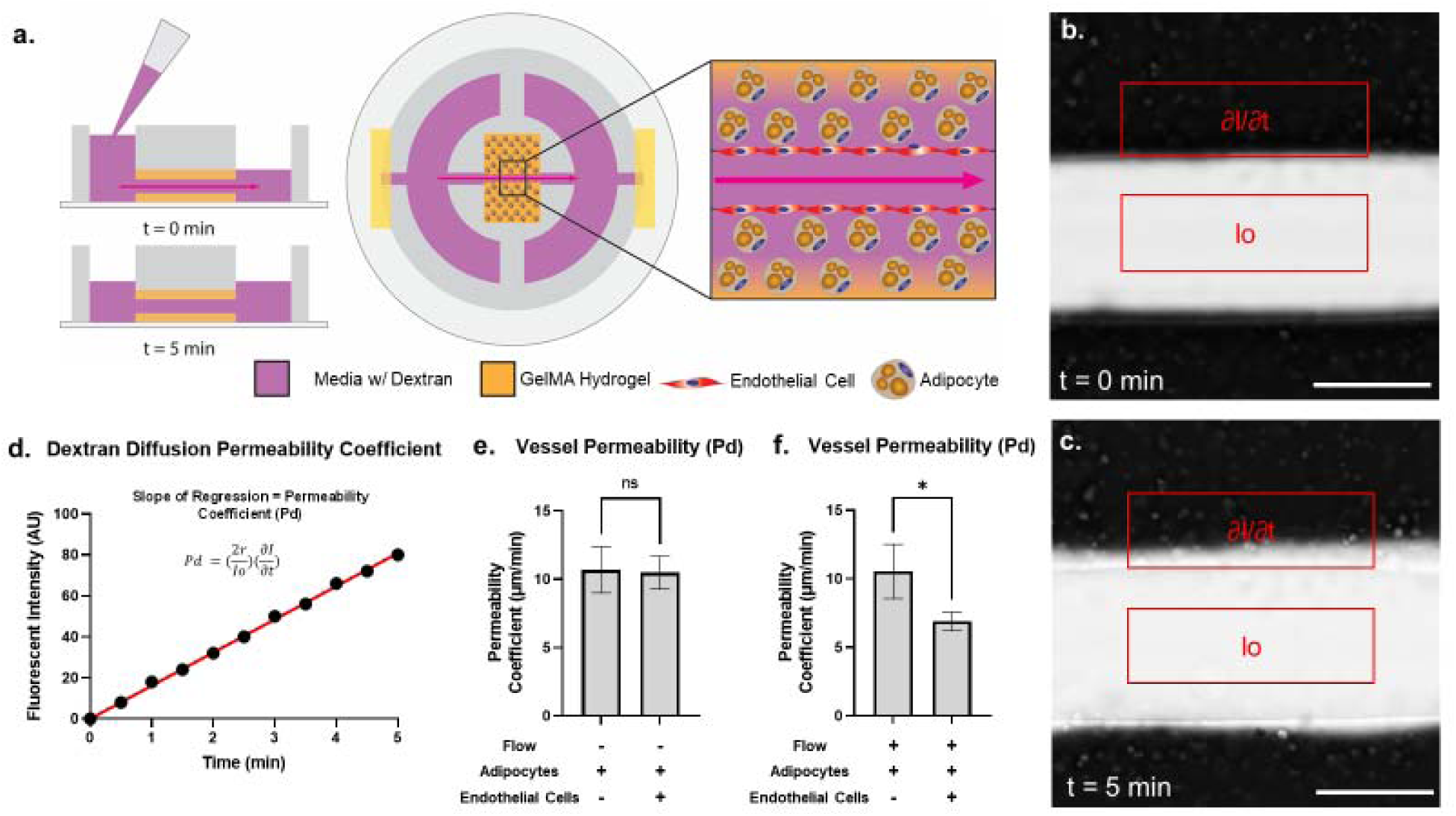
Vascular Permeability within µAT Platforms. (a) Experimental workflow for characterizing vascular permeability within µAT platforms. **(b,c)** Representative fluorescent images of dextran perfusion through the µAT platforms at **(b)** t = 0 minutes and **(c)** t = 5 minutes. Fluorescent intensity was obtained at the vessel region *(Io)* and gel region (∂*I/*∂*t)*. **(d)** Permeability coefficient *(Pd)* was calculated as a function of time and the slope of the regression line represents the rate of diffusion from the vessel into the surround hydrogel. **(e,f)** Quantitative analysis comparing **(e)** the presence of endothelial cells or **(f)** the presence of flow for each culturing condition. Data are presented as means ± SD. Comparisons between groups and statistical analysis were performed using two-tailed Student’s t-tests (*p < 0.05). Scale bar, 400 microns.

### Characterization of adipocyte function in µAT platforms

Adipocyte function within µAT platforms was evaluated by insulin-stimulated glucose uptake and lipolysis (figure 4(a) and 4(b)). No significant differences were detected in basal lipolysis for all conditions (figure 4(c)). However, insulin-stimulated glucose uptake was significantly increased when comparing µAT platforms cultured with endothelial cells with shear flow compared to the static adipocyte-only condition (figure 4(d)).

**Figure 4.**
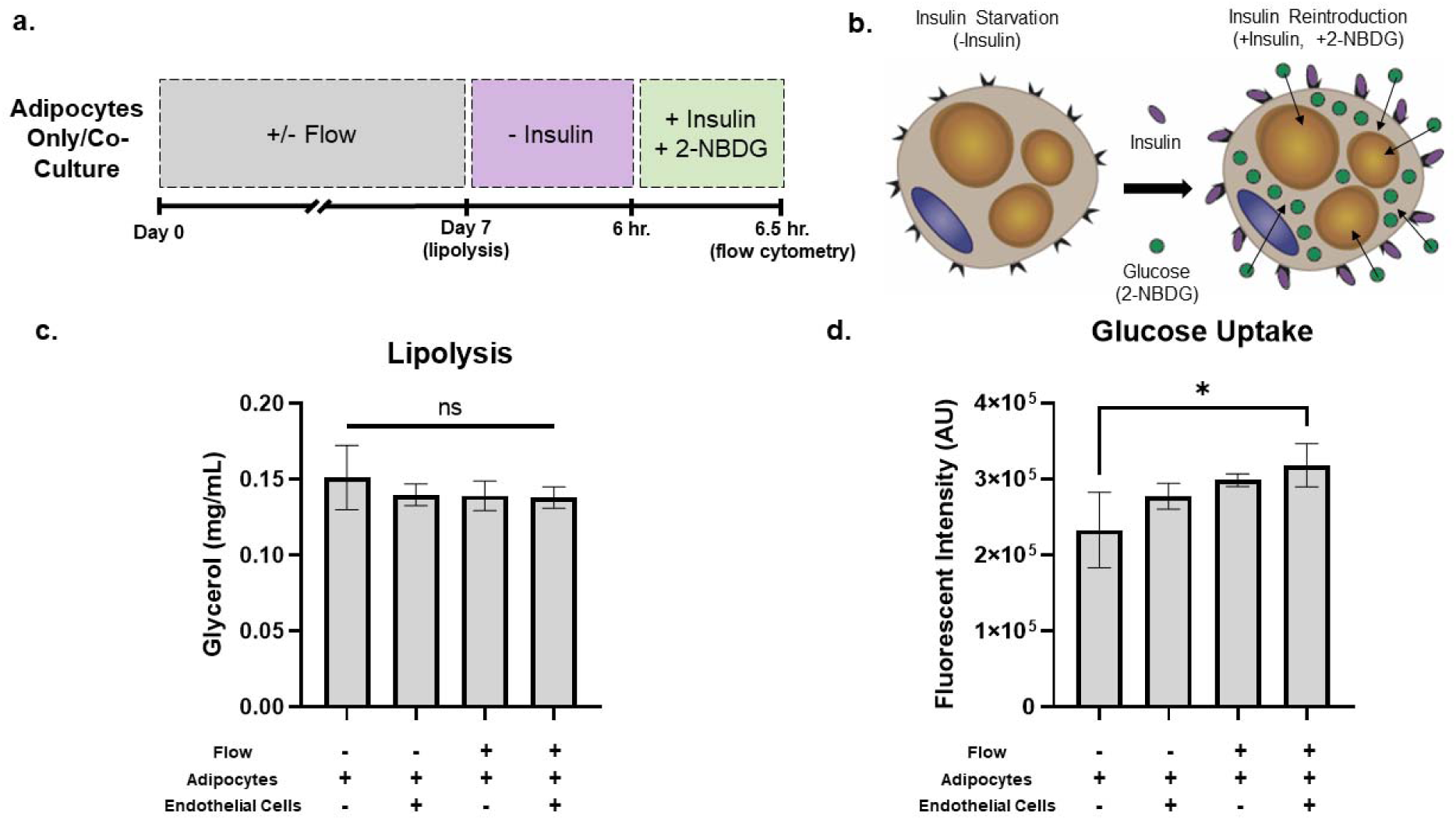
Characterization of Adipocyte Function in µAT Platforms. (a) Experimental workflow for collecting and analyzing adipocyte lipolysis and glucose uptake. **(b)** Visual representation of insulin induced uptake of fluorescent glucose analog (2-NBDG) in adipocytes. **(c,d)** Quantitative analysis of **(c)** lipolysis and **(d)** glucose uptake comparing the presence of endothelial cells or the presence of flow for each culturing condition. Data are presented as means ± SD. Comparisons between groups and statistical analysis were performed using one-way ANOVA (*p < 0.05).

### Vessel permeability within µAT platforms treated with forskolin

To determine if short-term forskolin exposure alters vascular permeability, on day 3, we exposed the µAT platforms to 10 µM forskolin for 4 hours prior to permeability assays. There was a significant decrease in vessel permeability in µAT platforms exposed to forskolin compared to the untreated group (figure 5(b)). Together these results suggest that the presence of shear flow decreases vascular permeability, and this decrease in permeability is further enhanced when exposed to forskolin (figure 5(c)).

**Figure 5.**
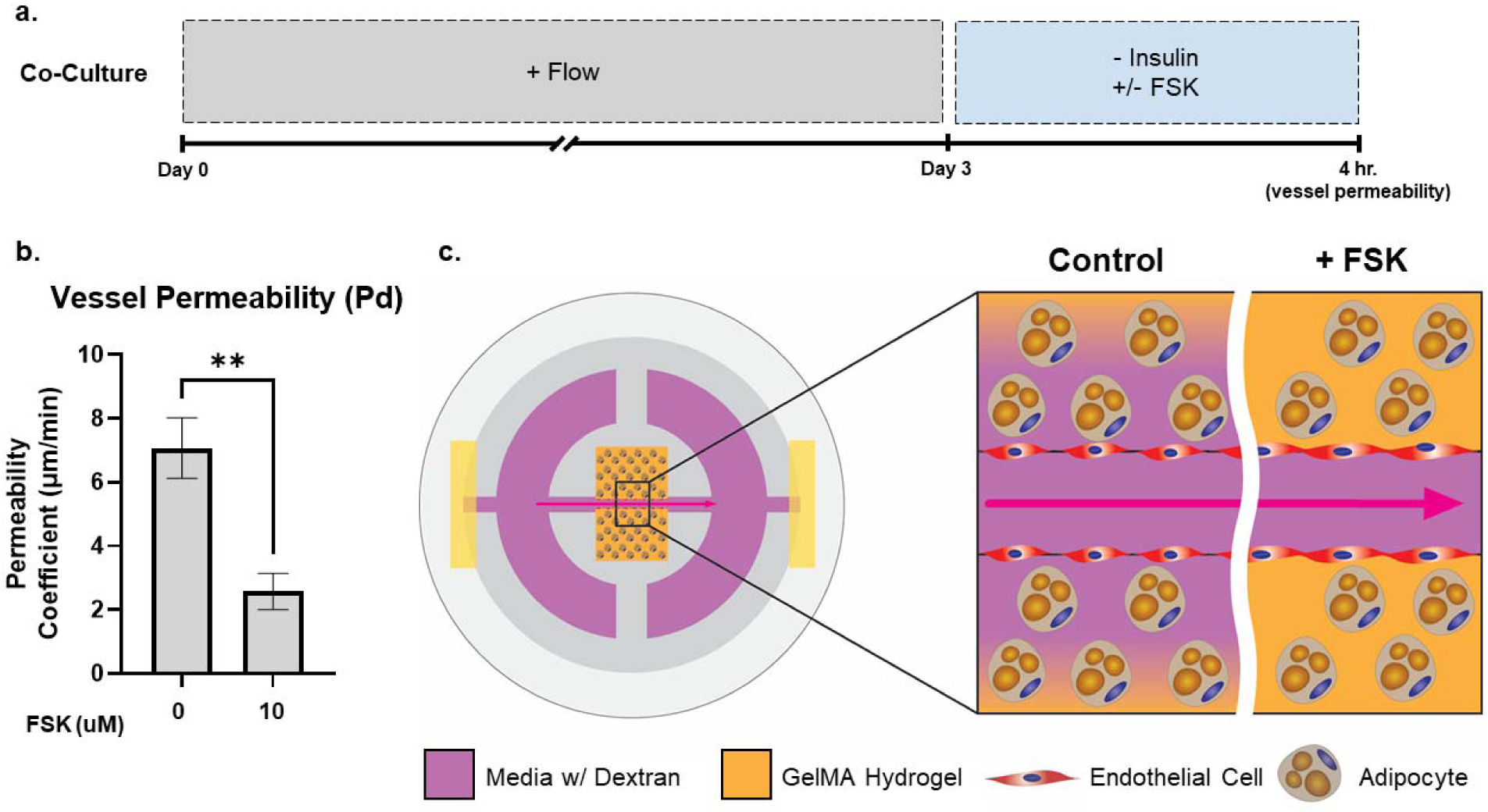
Vascular Permeability within µAT Platforms treated with Forskolin. (a) Experimental workflow for forskolin treatment. **(b)** Quantitative analysis of µAT platforms treatment with forskolin. **(c)** Visual representation of increased endothelial barrier function after treatment with forskolin. Data are presented as means ± SD. Comparisons between groups and statistical analysis were performed using two-tailed Student’s t-tests (*p < 0.05).

### Characterization of adipocyte function in µAT platforms treated with forskolin

To determine if forskolin exposure affected adipocyte function, we measured insulin-stimulated glucose uptake and lipolysis in µAT platforms (figure 6(a)). On day 7, µAT platforms were exposed to 10, 25, and 100 µM forskolin for 6 hours to determine if there is a concentration dependent effect on adipocyte function. Lipolysis remained unchanged after exposure µAT platforms to 10 µM forskolin. However, lipolysis was significantly increased when µAT platforms were exposed to 25 µM and 100 µM forskolin (figure 6(b)). For insulin-stimulated glucose uptake, no significant changes were found for any forskolin concentration (figure 6(c)). Together these results suggest adipocytes within µAT platforms respond to forskolin in a concentration-dependent manner, whereby lipolysis products are released into the vessel, modeling a physiologically relevant process.

**Figure 6.**
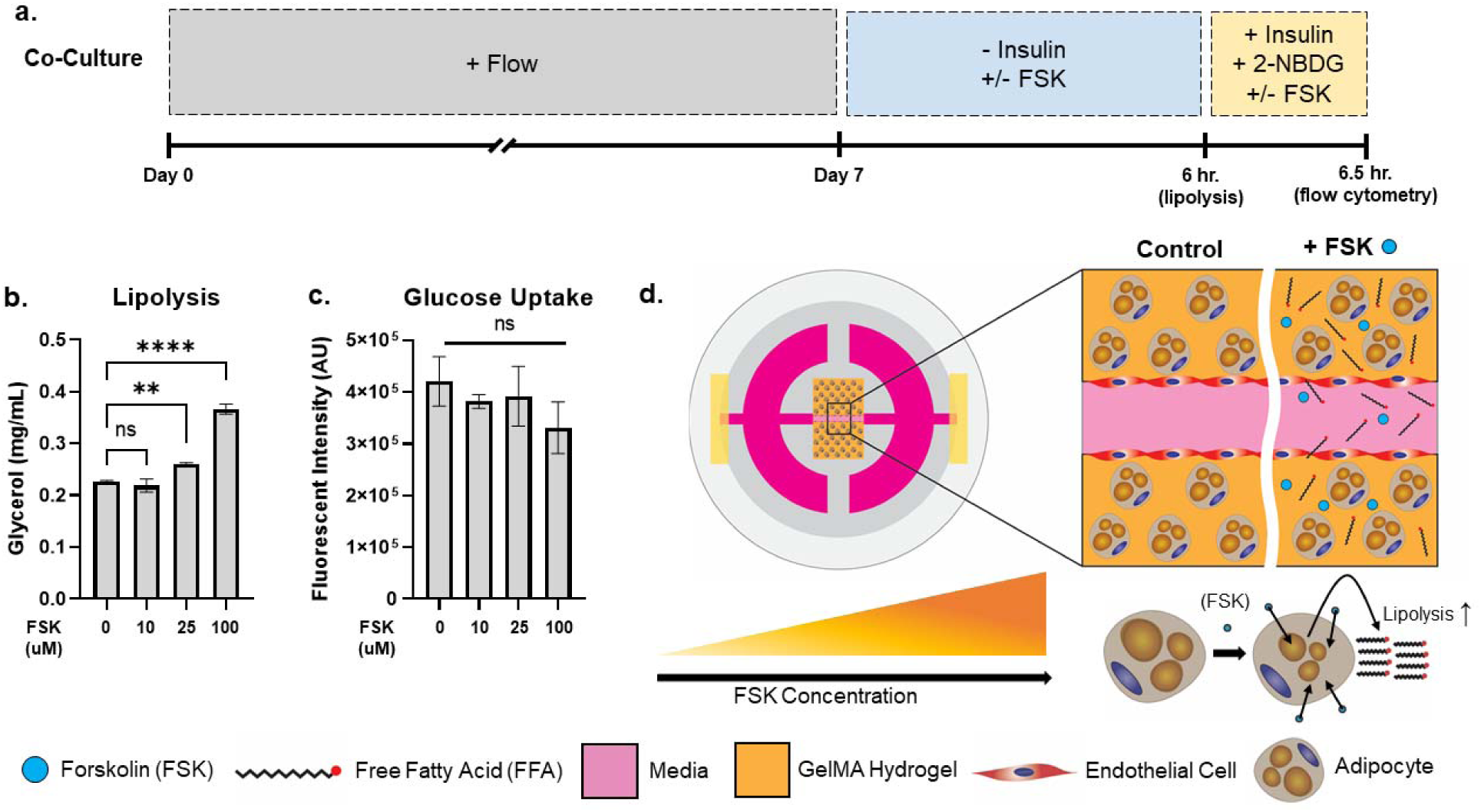
**Characterization of Adipocyte Function in µAT Platforms treated with Forskolin**. **(a)** Experimental workflow of forskolin treatment and collection of adipocyte lipolysis and glucose uptake. **(b,c)** Quantitative analysis of adipocyte **(b)** lipolysis and **(c)** glucose uptake using varying concentration of forskolin. **(d)** Visual representation of increased adipocyte lipolysis after treatment with forskolin. Data are presented as means ± SD. Comparisons between groups and statistical analysis were performed using one-way ANOVA (**p < 0.01, ****p < 0.0001).

## Discussion

In this study, we successfully fabricated µAT platforms (figure 1) and validated physiologically accurate shear flow (>3 dynes/cm^2^) using computational fluid dynamics (figure S1-S6)[24]. We confirmed a homogenous dispersion of adipocytes surrounding an open and perfusable vessel structure within the µAT bulk hydrogel region (figure 2(b) – 2(f)). When comparing static conditions, with and without endothelial cells lining the vessel, we did not observe a difference in vessel permeability. It is expected that dextran diffusion is slower in nanoporous GelMA hydrogels[31] compared to the more commonly employed microporous collagen hydrogels[32], which could mask the initial, early diffusion rate out of the vessel. However, the addition of shear flow caused a significant decrease in vessel permeability (figure 3(f)). In vitro endothelial cell monolayers or vessel structures exposed to shear flow are known to respond with increased endothelial cell alignment and improved cell-cell interactions to regulate permeability[26–30]. While changes in shear flow induced vascular permeability have been well characterized for other cell/tissue systems, little is known about how shear flow affects adipocyte function when endothelial cells or the vasculature are present.

Key adipocyte functional readouts were assessed through lipolysis and insulin-stimulated glucose uptake. Basal lipolysis was not affected by the presence of shear flow through the vessel or by the presence of endothelial cells (figure 4(c)), this matches our previous work where static co-cultures also did not have altered basal lipolysis[12]. However, insulin-stimulated glucose uptake was significantly increased in µAT platforms cultured with endothelial cells in the presence of shear flow compared to the adipocyte-only, static group (figure 4(d)). These differences in insulin-stimulated glucose uptake appear to be a synergistic effect of co-culture and shear flow, as neither flow nor endothelial cells alone with adipocytes yielded significant glucose uptake levels compared to adipocyte-only static control. Nitric oxide (NO) production has been shown in 3T3-L1 mouse adipocytes and HUVECs, and NO production increases in HUVECs under shear flow conditions[32,33]. Additionally, it has been reported that elevated NO levels increase the glucose uptake response through insulin-independent glucose transporter type 4 (GLUT4) translocation to the cell membrane in 3T3-L1 mouse adipocytes[32]. Therefore, the increased insulin-stimulated glucose uptake observed could be the result of increased NO production by endothelial cells experiencing shear flow within the vessel.

To show the µAT platform potential as a drug screening platform, the µAT platforms were exposed to the pharmacological compound forskolin. Forskolin is an adenyl cyclase agonist targeting similar downstream signaling pathways as β-adrenergic agonists, such as norepinephrine, without interacting with adrenergic receptors, which increase lipolysis and alter vascular permeability in a temporal and concentration dependent manner[34,35]. Short-term (1-2 hours) exposure to forskolin has been found to decrease permeability of an endothelial cell monolayer in a transwell assay and has been shown to increase lipolysis in 3T3-L1 mouse adipocytes[36,37]. Consistent with these previous findings, we found short-term (4-6 hours) forskolin exposure led to a decrease in vessel permeability (figure 5(b)), and a concentration dependent increase in lipolysis (figure 6(b)). Insulin-stimulated glucose uptake was not sensitive to any forskolin concentration (figure 6(c)). Forskolin has been shown to decrease insulin-stimulated glucose uptake in 3T3-L1 mouse adipocytes through the dissociation of the mammalian target of rapamycin (mTOR) complex, a downstream target of insulin signaling[38]. While our results do not show the same findings in the conditions shown, there is a concentration dependent decreasing trend in glucose uptake consistent with previous findings in insulin sensitive muscle cells[39], although not statistically significant.

While several MPS platforms have been used to model angiogenesis, tumor metastasis, and immune cell migration[6,19–23], little has been done to specifically model AT functions. Previously, one MPS platform of AT has used 2D adipocyte monolayers separated by a porous membrane allowing for enhanced nutrient and waste exchange[40]. The permeable membrane mimics the endothelium as a physical porous barrier, shielding adipocytes from high shear stresses during perfusion. However, this platform lacked critical interactions with an extracellular matrix and endothelial cell crosstalk. These limitations were then partly addressed by embedding adipocytes within 3D hydrogels[41,42], yet still lack the critical crosstalk between the adipocytes and their neighboring vasculature[8,10,12,15].

MPS platforms modeling AT with vascular components currently exist, with similar fabrication methods (PDMS device, subtractive molding for a vessel and bulk hydrogel encapsulating adipocytes) and some similar functional aspects (permeability, media sampling, visualization by confocal microscopy) as the µAT platform described here[43,44]. However, these AT MPS platforms have a much larger footprint and require 1-2 syringe pumps per platform, severely limiting their throughput. The µAT platform described in this study addresses this limitation via integration with traditional flat bottom 24 well plates without the need for pumps. Additionally, these platforms often rely on indirect measurements of adipocyte function, such as lipogenesis (the accumulation of lipids within adipocytes) by measuring adipocyte or lipid droplet size. In this study, adipocyte function can be measured more directly via insulin stimulated glucose uptake and lipolysis. Unlike other platforms and 3D models, these functions are performed by media perfusion and sampling via a central blood vessel, lending itself to improved physiological relevance. Moreover, the µAT platforms can achieve physiological shear flow without these external pumps by gravity-driven perfusion, thus considerably reducing the chances of air bubble formation and contamination during media changes. Therefore, the design considerations implemented in this µAT platform significantly increased the experimental throughput to allow for more facile screening of multiple factors at once.

## Conclusions

The µAT platform allows for the testing of key AT functions (lipolysis, glucose uptake) via a physiologically responsive vessel in a multi-throughput format for scaling up to therapeutic screening. Simple integration into multi-throughput workflows makes the µAT platform an ideal candidate for therapeutic testing platforms and personalized medicine by modeling essential aspects of AT and AT-related diseases (figure 7). Further studies include the incorporation of various obesogenic cues and different endothelial cell sources to model diseases within AT, such as lipedema and lymphedema, which remain poorly understood.

**Figure 7.**
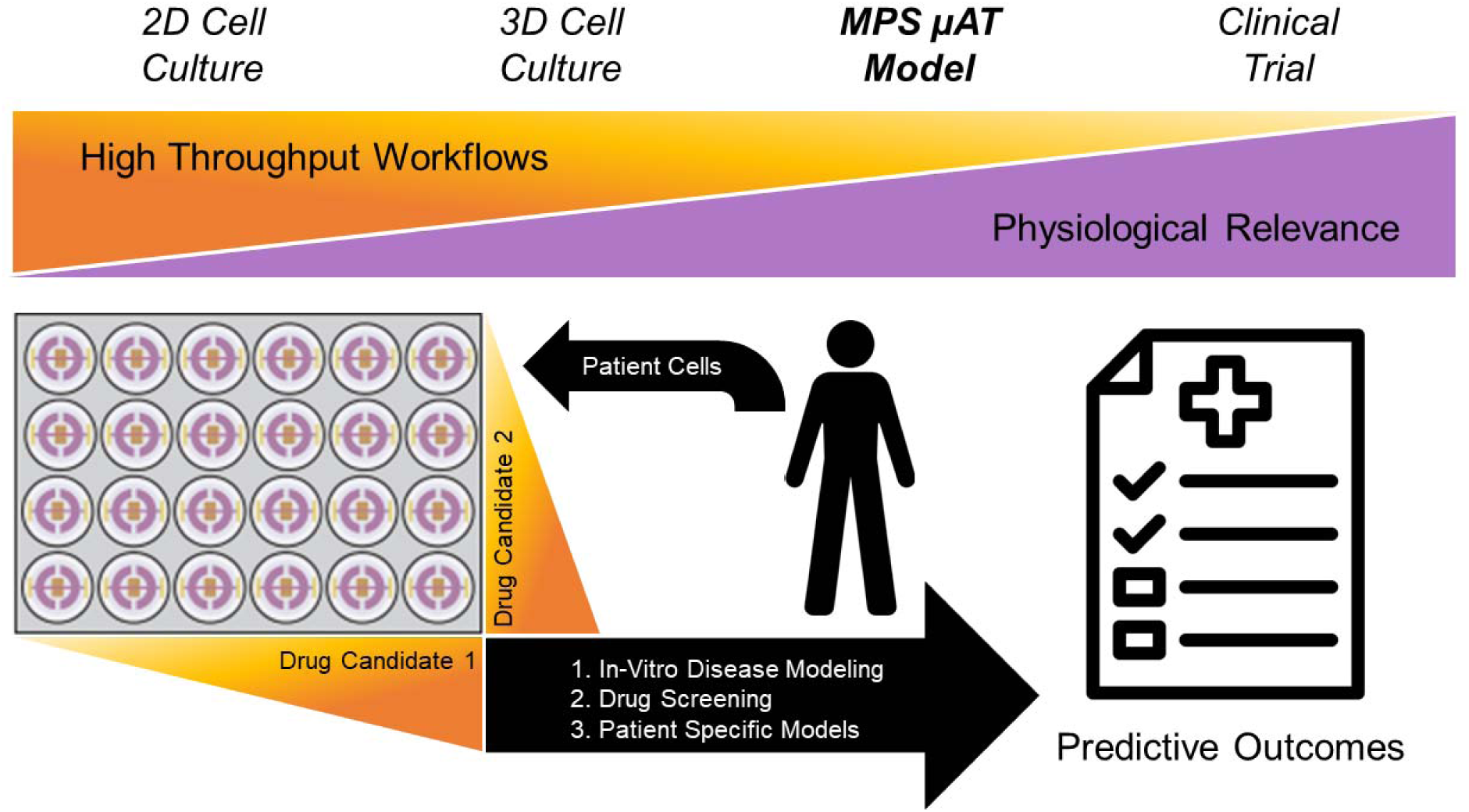
µAT Platforms Integration into Multi-Throughput Workflows. The µAT platform is an ideal candidate for therapeutic testing platforms and personalized medicine by modeling essential aspects of AT and AT-related diseases.

## Author contributions

M.S. performed all experiments and data analysis. M.S. and E.B. designed all experiments, interpreted the results, and wrote the manuscript. Both authors reviewed the manuscript.

## Acknowledgments

The authors would like to acknowledge funding support from the NIH NIDDK Diabetic Complications Consortium DK07616 and DK115255 (to E.B.), NSF CAREER 2045517 (to E.B.) for their financial support toward the project.

---

Key resources table.

Key resources table
| REAGENT or RESOURCE | SOURCE | IDENTIFIER |
| --- | --- | --- |
| **Antibodies** |  |  |
| Anti-CD31 mouse monoclonal | Invitrogen | Cat# PIMA513188, RRID: AB_10982120 |
| Alexa Fluor™ 647 conjugated Rabbit anti-Mouse IgG (H+L) | Invitrogen | Cat# A21239, RRID: AB_2535808 |
| **Bacterial and virus strains** |  |  |
| n/a |  |  |
| **Biological samples** |  |  |
| n/a |  |  |
| **Chemicals, peptides, and recombinant proteins** |  |  |
| Low glucose Dulbecco's Modified Eagle's medium (DMEM) | Corning | 10014CM |
| Fetal bovine serum (FBS) | Sigma-Aldrich | 12306C-500ML |
| Penicillin/streptomycin (P/S) | Cytiva | SV30010 |
| Dulbecco's Modified Eagle Medium/Nutrient Mixture F-12 (DMEM/F12) | Corning | 10092CM |
| Rosiglitazone | Cayman Chemical | 71740100, CAS: 122320-73-4 |
| 3-isobutyl-1-methylxanthine | Thermo Scientific | AC228420010, CAS: 28822-58-4 |
| Dexamethasone | Thermo Scientific | AC230302500, CAS: 50-02-2 |
| D-(+)-Biotin | Thermo Scientific | AAA1420703, CAS: 58-85-5 |
| D-calcium pantothenate | TCI America | P0012100G, CAS: 137-08-6 |
| Recombinant human insulin | Sigma Aldrich | 50-165-6853, CAS: 11061-68-0 |
| EGM-2 Endothelial Cell Growth Medium-2 BulletKit | Lonza | CC-3162 |
| Polydimethylsiloxane (PDMS) – Sylgard 184 | Ellsworth | DC2065622 |
| Isopropanol | Fisher | A4154, CAS: 67-63-0 |
| Gelatin | Sigma-Aldrich | G2500, CAS: 9000-70-8 |
| Sodium bicarbonate | Fisher | S233-500, CAS: 144-55-8 |
| Methacrylic anhydride (94%) | Alfa-Aesar | L14357, CAS: 760-93-0 |
| phosphate buffer saline | Fisher | BP661-50 |
| Formaldehyde | Thermo Scientific | 28908 |
| Bovine serum albumin (BSA) | Fisher | BP9703100, CAS: 9048-46-8 |
| BODIPY 493/503 (fluorescent lipid stain) | Invitrogen | D3922, CAS: 121207-31-6 |
| Hoechst 33342 (fluorescent nuclear stain) | Thermo Scientific | 62249, CAS: 875756-97-1 |
| Dextran, Cascade Blue™, 10,000 MW, Anionic, Lysine Fixable | Invitrogen | D1976 |
| 2-Nitrobenz Oxadiazol Amino Deoxyglucose (2-NBDG) | Invitrogen | N13195, CAS: 186689-07-6 |
| DRAQ5 (fluorescent nuclear stain) | Thermo Scientific | 62254, CAS: 254098-36-7 |
| Collagenase type I | Gibco | 17100017 |
| Forskolin | Sigma Aldrich | F6886, CAS: 66575-29-9 |
| Critical commercial assays |  |  |
| Sekure Triglyceride Reagent | Sekisui Diagnostics | NC1803782 |
| Deposited data |  |  |
| n/a |  |  |
| Experimental models: Cell lines |  |  |
| Human bone marrow (to derive mesenchymal stem cells) | Lonza | 1M-105 |
| Human umbilical vein endothelial cells | Lonza | C2519A |
| Experimental models: Organisms/strains |  |  |
| n/a |  |  |
| Oligonucleotides |  |  |
| n/a |  |  |
| Recombinant DNA |  |  |
| n/a |  |  |
| Software and algorithms |  |  |
| OnShape | OnShape | Free plan |
| ImageJ | NIH | FIJI package |
| ANSYS | ANSYS Inc. | Student License |
| GraphPad Prism | GraphPad Software Inc. | Prism 9 |
| Other |  |  |
| 3D printer | Stratasys | Objet Connex2 350 |
| 15 mm round glass coverslips | Electron Microscopy Sciences | 7222801 |
| Plasma asher | Plasma Etch | PE-25 |
| Dialysis tubes | Fisherbrand | 21-152-14 |
| 0.22 µm filter | Fisher | SCGVU05RE |
| 400-micron stainless steel rod | Myotech | 7100528 |
| 405 nm lamp (20W lamp) | Amazon | B085NMPW7B |
| Vacuum grease | Corning | 2966k52 |
| 2D Rocking Platform | Denville | S2036 |
| Inverted laser scanning confocal microscope | Olympus | FluoView FV1200 |
| 10X objective | Olympus | UPLXAPO10X |
| Plate reader | TECAN | Infinite M200 Pro |
| Inverted spinning disk microscope | Olympus | IX-81 |
| Flow cytometer | BD Biosciences | Accuri C6 |

## Resource availability

### Lead contact

Further information and requests for resources and reagents should be directed to and will be fulfilled by the lead contact, Evangelia Bellas.

### Materials availability

This study did not generate new unique reagents.

## Data and code availability

All data needed to evaluate the conclusions in the paper are present in the paper and the supplemental information. Any additional information required to reanalyze the data reported in this paper is available from the lead contact upon request.

## Experimental model and study participant details

### Cell culture

Primary cell cultures were employed in this study. Human mesenchymal stem cells were isolated from human bone marrow purchased from Lonza: Catalog #: 1M-105, Lot #: 0000845857, Age: 23, Sex: Male, Race: Hispanic. Human umbilical vein endothelial cells were purchased from Lonza: Catalog #: C2519A, Lot #: 0000631340, Age: newborn, Sex: Male/female mix, Race: Caucasian/African American.

Human bone marrow-derived mesenchymal stem cells (hMSCs) were isolated from healthy adult male bone marrow, purchased from Lonza Walkersville Inc. (Walkersville, MD, Cat# 1M-105). HMSCs were expanded in low glucose Dulbecco’s Modified Eagle’s medium (DMEM, Corning) supplemented with 10% (v/v) fetal bovine serum (FBS, Sigma-Aldrich) and 1% (v/v) penicillin/streptomycin (P/S, Cytiva) in 2D until confluency. Adipogenic induction media was added when hMSCs reached confluency at passage 4. The adipogenic induction media was composed of Dulbecco’s Modified Eagle Medium/Nutrient Mixture F-12 (DMEM/F12, Corning) supplemented with 3% (v/v) FBS, 1% (v/v) P/S, 2 µM rosiglitazone (Cayman Chemical), 500 µM 3-isobutyl-1-methylxanthine (Sigma-Aldrich), 1 µM dexamethasone (Acros Organics), 33 µM biotin (Alfa Aesar), 17 µM D-calcium pantothenate (TCI America), and 20 nM of insulin (Sigma-Aldrich). The hMSCs were differentiated for 7 days prior to encapsulation within bulk hydrogel region.

Human umbilical vein endothelial cells (HUVECs) were purchased from Lonza Walkersville Inc. (Walkersville, MD, Cat# C2519A) and expanded to passage 8 endothelial growth media (EGM-2, Lonza). Endothelial cells were cultured to ∼80% confluency with media changes every 2-3 days until seeded in µAT platforms. All cultures were maintained at 37°C and 5% CO_2_.

## Method details

### Template fabrication and preparation

The µAT platforms were fabricated with polydimethylsiloxane (PDMS, Sylgard 184, Ellsworth) using a 3D-printed reverse mold template. The reverse mold template was custom designed using CAD software (OnShape) and printed using the Objet Connex2 350 3D printer (Stratasys) with Vero glossy finish photo resin. A PDMS prepolymer, 10:1 (w/w) ratio of base polymer to curing agent, was cast against the reverse mold template, and thermally cured at 100°C for 45 minutes. The cured µAT platforms were separated from the reverse mold template, cut to shape, and soaked in 100% isopropanol (Fisher) overnight to remove uncrosslinked PDMS oligomers. The µAT platforms were placed at 60°C for 4 hours before plasma treatment to evaporate residual isopropanol. µAT platforms and 15 mm round glass coverslips (Electron Microscopy Sciences) were treated with air plasma (Plasma Etch, PE-25, 75W, 2 min) for covalent bonding and placed at 60°C overnight to enhance the adhesion of the PDMS to glass. µAT platforms were sterilized under ultraviolet irradiation for 15 minutes before cell seeding.

### Gelatin methacryloyl (GelMA) preparation

10 grams of gelatin (Sigma-Aldrich, type A, 300 bloom) were dissolved in 100 mL of 0.25 M sodium bicarbonate (Fisher) buffer at 50°C on a stirring hot plate until completely dissolved. Once dissolved, 2.5 mL (2.5% v/v) Methacrylic anhydride (Alfa Aesar, 94%) was added dropwise at a flow rate of 0.6 mL/minute to the gelatin solution and allowed to react for 1 hour. During the reaction, the pH was adjusted to 9 and protected from light. The gelatin methacryloyl (GelMA) solution was diluted at 1:4 in prewarmed 1X phosphate buffer saline (PBS) and placed in dialysis tubes (Fisherbrand, 12-14 kDa MWCO). The GelMA solution was dialyzed for 3 days against 3L of 1X PBS followed by 3 days against 3L of deionized water. All dialysis was performed at 55°C on a stirring hot plate and protected from light. After dialysis, the GelMA solution was sterile and filtered through a 0.22 µm filter (Fisher), aliquoted into sterile tubes, snap-frozen, and lyophilized. The lyophilized GelMA was stored at –80°C until use.

### Assembly of µAT platform

The µAT platforms were designed to be compatible with the traditional flat bottom 24 well plates (figure 1(a)). The platforms incorporate two open top media reservoirs, one central microchannel, and one closed top hydrogel reservoir in the center of the devices (figure 1(b) and S7), modeling a blood vessel within AT. The microchannel connects the two media reservoirs, allowing media perfusion across the microfluidic device. The µAT platforms were fabricated by inserting a 400-micron stainless steel rod (Myotech) through the microchannel guides before filling it with unpolymerized GelMA precursor (figure 1(c)). Adipocytes were resuspended in a 15% (w/v) gelatin methacryloyl (GelMA) precursor solution for a final concentration of 4 million cells/mL. The adipocyte-GelMA solution was loaded into the µAT platforms and polymerized with 405 nM (Amazon, 42 mW) light for 2 minutes (figure 1(d)). The stainless-steel rod was removed, forming a hollow microchannel via subtractive molding. The open ends of the microchannels were sealed using bioinert vacuum grease (Corning) to prevent leakages (figure 1(f)). Media was added to the media reservoirs and maintained at 37°C for 30 minutes, after which the endothelial cells were seeded. Passage 8 endothelial cells were resuspended in EGM-2 media at 10 million cells/mL. 60 µL of the endothelial cell suspension was pipetted into media reservoirs and allowed to attach to the microchannel for 1 hour. This process was repeated twice, once upright and once inverted, to allow endothelial cells to attach and cover the top and bottom of the microchannel (figure 1(e) and 1(f)). Media reservoirs were then aspirated, washed 3 times with 1X PBS, and filled with 240 µL fresh media. Day 1 is counted as the day the devices were fabricated and seeded.

### µAT platform culture

The µAT platforms were cultured with a 1:1 (v/v) ratio of EGM-2 and adipogenic maintenance media (adipogenic induction media without rosiglitazone and 3-isobutyl-1-methylxanthine). Media was exchanged daily. The µAT platforms were maintained in static or flow conditions (Biorocker 2D Rocking Platform, Denville, S2036, 30° maximum rocking angle, 0.5 Hz rocking speed) for 7 days (figure 1(i)).

### Visualization by immunofluorescence and cell staining

After 7 days, µAT platforms were fixed in 4% (w/v) formaldehyde (Thermo Scientific) in 1X PBS for 1 hour, washed 3 times with 1X PBS, then blocked with 3% (w/v) bovine serum albumin (BSA, Fisher Scientific) in 1X PBS overnight. The µAT platforms were incubated overnight with an mouse anti-human CD31 primary antibody (Invitrogen, Cat# MA5-13188, RRID:AB 10982120, 1:200 dilution), washed 3 times with 1X PBS, and incubated overnight with a donkey anti-mouse IgG Alexa Fluor® 647 secondary antibody (Invitrogen, Cat# A-31571, RRID:AB_162542, 1:1000 dilution). The µAT platforms were co-stained with BODIPY 493/503 (Invitrogen, 1:200 dilution) and Hoechst 33342 (Thermo Fisher, 1:1000 dilution) for 1 hour for lipid and nuclei visualization, respectively. All fluorescent and immunofluorescent stain incubation was performed on a plate rocker at room temperature. Samples were stored in 1X PBS at room temperature until imaged.

### Fluorescent confocal Imaging and image analysis

Images were acquired with an inverted laser scanning confocal microscope (Olympus FluoView FV1200) using a 10X objective (UPLXAPO10X, 0.4 NA, Olympus). Images were imported to ImageJ for processing. Z-stacks (thickness ∼ 400 µm, step-size: 5 µm, 10 µs/pixel, 1024×1024) were compressed, and maximum intensity projections were used to visualize the organization within the platforms.

### Vascular permeability assay

After 3 days, a vascular permeability assay was performed by perfusing fluorescent dextran (Invitrogen, Cascade Blue™, 10,000 MW, 12.5 µg/mL) containing media at an average hydrostatic pressure difference of ∼ 0.16 cm H_2_O. Fluorescent intensity was obtained at the vessel region *(Io)* and hydrogel region (∂*I/*∂*t)* at every time point, and the permeability coefficient *(Pd)* was calculated using equation 1 where *r*, *Io*, ∂*I/*∂*t* represent the radius of the vessel, fluorescent intensity of the vessel region, and the fluorescent intensity flux of the hydrogel region over time (figure 3(b) and 3(c)).

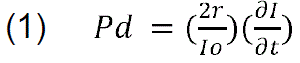

Images were acquired in real-time with an inverted spinning disk confocal (Olympus IX-81) using a 10X objective (UPLXAPO10X, 0.4 NA, Olympus) with 5 second exposure time, 30 seconds per image for 5 minutes. The fluorescent intensity of the hydrogel region (∂*I/*∂*t)* was normalized to zero at t = 0, and changes in hydrogel intensity over time were plotted. The permeability coefficient *(Pd)* was determined using a linear fit regression with the y-intercept set to 0 to obtain the slope (figure 3(d)).

### Lipolysis assays

After 7 days, media was sampled from the media reservoirs and banked at –80°C until further use. Glycerol content, a byproduct of triglyceride hydrolysis, was measured using Sekure Triglyceride Reagent (Sekisui Diagnostics) according to the manufacturer protocol. Absorbance at 520 nm was quantified using an Infinite M200 Pro plate reader (TECAN).

### Insulin-stimulated glucose uptake

After 7 days, insulin-stimulated glucose uptake was performed. µAT platforms were insulin starved for 6 hours followed by insulin reintroduction using media containing 20 µM fluorescent glucose analog, 2-Nitrobenz Oxadiazol Amino Deoxyglucose (2-NBDG, Invitrogen, Cat# N13195) for 30 minutes (figure 4(a)). µAT platforms were washed 3 times in 1X PBS and fixed in 4% (w/v) formaldehyde in 1X PBS for 1 hour. After fixation, µAT platforms were incubated with nuclear stain, DRAQ5 (Thermo Fisher, 1:1000 dilution) in 1X PBS for 1 hour followed by enzymatic digestion in collagenase type I (35 mg/mL, Corning) for 30 minutes at 37°C. The digested cell suspension was resuspended in 1X PBS to a final volume of 300 µL, and fluorescent intensity was quantified using flow cytometry (Accuri C6, BD Biosciences). Hydrogel and cellular debris were removed by excluding events with less than 10^3 fluorescent intensity in the DRAQ5 channel (FL4-A, 647 nm excitation) and deleting events less with less than 2,000,000 FSC-H and 500,000 SSC-H. Adipocyte populations were identified within scatter FSC-A vs. SSC-A plots, and mean fluorescent intensities (FL1-A, 488 nm excitation) were recorded.

### Forskolin treatment

On day 3, a vascular permeability assay was performed by exposing µAT platforms to forskolin (Sigma) for 4 hours (figure 5(b)). For vascular permeability assays, fluorescent dextran containing media +/− forskolin was used, and images were acquired as described above. For lipolysis assay, on day 7, media was replaced with media containing +/− forskolin and incubated for 6 hours before collecting media (figure 6(a)). Spent media was collected and banked at – 80°C until further use. For insulin-stimulated glucose uptake assays, on day 7, µAT platforms were insulin starved for 6 hours +/− forskolin followed by insulin reintroduction +/− forskolin using 2-NBDG as described above (figure 6(a)).

### Flow calculations and computational fluid dynamics (CFD)

Using the properties of water at 37°C, wall shear stress (τ) at the maximum rocking angle (30°) was calculated to be approximately 0.487 Pascals, or 4.87 dynes/cm^2^ using equation (2) where *r*, *p*, *g*, α represent the vessel radius, density of water, gravity, and the maximum angle of the biorocker[24].

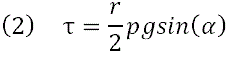

Water at 37°C was used in the mathematical calculations and ANSYS simulations since it is a commonly used default material. Mathematical calculations confirmed a physiologically accurate (>3 dynes/cm^2^) wall shear stress experienced on the vessel within the µAT platform[24]. The volumetric flow rate *(Q)* at the maximum rocking angle was calculated to be 3.96*10^-3^ cm^3^/second using equation (3) where *r*, *u*, *t* represent the vessel radius, dynamic viscosity of water, and wall shear stress previously calculated in equation (2)[24].

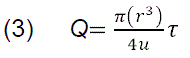

The pressure drop (D*P)* at the maximum rocking angle was calculated to be 24.36 Pascals using the Hagen – Poiseuille equation (4) for Poiseuille flow in a pipe where *u*, *L*, *D*, *Q* are the dynamic viscosity of water, vessel length, vessel diameter, and volumetric flow rate calculated in equation (3).

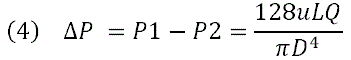

To ensure the volumetric flow rate *(Q)* at the maximum rocking angle considered hydraulic resistance, or fluids resistance to flow, we used equations (5,6) where *u*, *L*, *r*, Δ*P* represent the viscosity of water, vessel length, vessel radius, and pressure drop calculated in equation (4)[25].

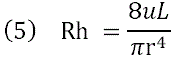

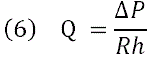

Additionally, we further validated our shear stress (τ) calculation by converting the pressure drop to wall shear stress using equation (7), where Δ*P*, *D*, *L* represent the pressure drop calculated in equation (4), vessel diameter, and vessel length.

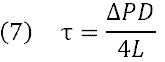

Next, we calculated the average fluid velocity *(V)* at the maximum rocking angle to be 3.15 cm/s using equation (8), where *Q* and *r* represent the volumetric flow rate calculated in equation (3) and vessel radius.

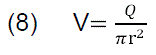

Lastly, laminar flow conditions at the maximum rocking angle were verified (16.19 << 2300) using the Reynolds number equation (9), where *p*, *V*, *D*, *u* represent density of water, fluid velocity, vessel diameter, and dynamic viscosity of water.

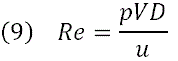

Ansys Fluid Flow (CFX) was used for computational fluid dynamics to confirm the mathematical calculation for wall shear stress calculated above. First, a geometry was created consisting of a 5 mm long cylinder with a diameter of 400 microns (figure S1). Next, a linear mesh was created using a 0.1 mm mesh size and was assigned an inlet, outlet, and wall (figure S2 and S3). In the experimental setup, the solver control was set to perform 100 iterations with a residual target (allowable error) to be 0.00001 and a steady-state analysis type was used. The average fluid velocity (V), calculated above, was used at the inlet of the cylinder, a relative pressure of 0 Pascals was used at the outlet of the cylinder, and no-slip wall conditions were used at the walls of the cylinder. In CFD-Post, after simulations have been completed, a 400-micron planar line was drawn in the y-direction across the diameter of the cylinder to plot the fluid velocity (V) and wall shear stress (τ) (figure S4-S6). ANSYS simulations approximate the wall shear stress to be 0.4854 – 0.4898 Pascals, consistent with the 0.487 pascals obtained through mathematical calculations (figure S6).

## Quantification and statistical analysis

GraphPad Prism 9 (GraphPad Software Inc.) software was used to perform all statistical analyses. All experiments were conducted with at least three biological replicates, and for each independent experiment, the averages of technical replicates were used. The distribution of each data set was analyzed, and the D’Agostino-Pearson test (α = 0.05) was performed to test for normality. Statistical comparisons between two experimental groups were performed using a two-tailed Student’s t-test. Comparisons among more groups were performed using one-way analysis of variance (ANOVA) with Tukey post hoc testing. All graphs are presented as mean ± standard deviations (SD) unless otherwise stated. Significance was determined according to the following p values: *p < 0.05, **p ≤ 0.01, ***p ≤ 0.001, and ****p ≤ 0.0001.

## References

[1] Choe S S, Huh J Y, Hwang I J, Kim J I and Kim J B 2016 Adipose tissue remodeling: Its role in energy metabolism and metabolic disorders *Front*. Endocrinol. (Lausanne). 7 1–16

[2] Ronti T, Lupattelli G and Mannarino E 2006 The endocrine function of adipose tissue: An update *Clin*. Endocrinol. (Oxf). 64 355–65

[3] Coelho M, Oliveira T and Fernandes R 2013 Biochemistry of adipose tissue: an endocrine organ. Arch. Med. Sci. 9 191–200

[4] Sengenès C, Bourlier V, Galitzky J, Zakaroff-Girard A, Lafontan M and Bouloumié A 2011 Cellular Remodeling during the Growth of the Adipose Tissue *Glob. Perspect*. Child. Obes. 183–90

[5] Longo M, Zatterale F, Naderi J, Parrillo L, Formisano P, Raciti G A, Beguinot F and Miele C 2019 Adipose tissue dysfunction as determinant of obesity-associated metabolic complications *Int*. J. Mol. Sci. 20

[6] Wu Q, Liu J, Wang X, Feng L, Wu J, Zhu X, Wen W and Gong X 2020 Organ-on-a-chip: Recent breakthroughs and future prospects *Biomed. Eng*. Online 19 1–19

[7] Chen F-M, Liu X, Polym P and Author S 2016 Advancing biomaterials of human origin for tissue engineering HHS Public Access Author manuscript Prog Polym Sci 53 86–168

[8] Bellas E, Marra K G and Kaplan D L 2013 Sustainable three-dimensional tissue model of human adipose tissue. Tissue Eng. Part C. Methods 19 745–54

[9] Anvari G and Bellas E 2021 Hypoxia induces stress fiber formation in adipocytes in the early stage of obesity *Sci*. Rep. 11 1–11

[10] Choi J H, Bellas E, Gimble J M, Vunjak-Novakovic G and Kaplan D L 2011 Lipolytic function of adipocyte/endothelial cocultures Tissue Eng. – Part A 17 1437–44

[11] Di Caprio N and Bellas E 2020 Adipose Tissue Pathology: Collagen Stiffness and Architecture Regulate Fibrotic Gene Expression in Engineered Adipose Tissue (Adv. Biosys. 6/2020) Adv. Biosyst. 4 2070061

[12] Hammel J H and Bellas E 2020 Endothelial cell crosstalk improves browning but hinders white adipocyte maturation in 3D engineered adipose tissue *Integr*. Biol. (Camb). 12 81–9

[13] Bellas E, Rollins A, Moreau J E, Lo T, Quinn K P, Fourligas N, Georgakoudi I, Leisk G G, Mazan M, Thane K E, Taeymans O, Hoffman A M, Kaplan D L and Kirker-Head C A 2015 Equine model for soft-tissue regeneration J. Biomed. Mater. Res. – Part B Appl. Biomater. 103 1217–27

[14] Bellas E, Lo T J, Fournier E P, Brown J E, Abbott R D, Gil E S, Marra K G, Rubin J P, Leisk G G and Kaplan D L 2015 Injectable Silk Foams for Soft Tissue Regeneration *Adv*. Healthc. Mater. 4 452–9

[15] Bellas E, Panilaitis B J B, Glettig D L, Kirker-Head C A, Yoo J J, Marra K G, Rubin J P and Kaplan D L 2013 Sustained volume retention in vivo with adipocyte and lipoaspirate seeded silk scaffolds Biomaterials 34 2960–8

[16] Ward A, Quinn K P, Bellas E, Georgakoudi I and Kaplan D L 2013 Noninvasive Metabolic Imaging of Engineered 3D Human Adipose Tissue in a Perfusion Bioreactor PLoS One 8 1–8

[17] Bellas E, Seiberg M, Garlick J and Kaplan D L 2012 In vitro 3D Full-Thickness Skin-Equivalent Tissue Model Using Silk and Collagen Biomaterials *Macromol*. Biosci. 12 1627–36

[18] Quinn K P, Bellas E, Fourligas N, Lee K, Kaplan D L and Georgakoudi I 2012 Characterization of metabolic changes associated with the functional development of 3D engineered tissues by non-invasive, dynamic measurement of individual cell redox ratios Biomaterials 33 5341–8

[19] Su C, Menon N V, Xu X, Teo Y R, Cao H, Dalan R, Tay C Y and Hou H W 2021 A novel human arterial wall-on-a-chip to study endothelial inflammation and vascular smooth muscle cell migration in early atherosclerosis Lab Chip 21 2359–71

[20] Xu Z, Li E, Guo Z, Yu R, Hao H, Xu Y, Sun Z, Li X, Lyu J and Wang Q 2016 Design and Construction of a Multi-Organ Microfluidic Chip Mimicking the in vivo Microenvironment of Lung Cancer Metastasis *ACS Appl*. Mater. Interfaces 8 25840–7

[21] Kim S, Lee H, Chung M and Jeon N L 2013 Engineering of functional, perfusable 3D microvascular networks on a chip Lab Chip 13 1489–500

[22] Margolis E A, Cleveland D S, Kong Y P, Beamish J A, Wang W Y, Baker B M and Putnam A J 2021 Stromal cell identity modulates vascular morphogenesis in a microvasculature-on-a-chip platform Lab Chip 21 1150–63

[23] Wang W Y, Lin D, Jarman E H, Polacheck W J and Baker B M 2020 Functional angiogenesis requires microenvironmental cues balancing endothelial cell migration and proliferation Lab Chip 20 1153–66

[24] Polacheck W J, Kutys M L, Tefft J B and Chen C S 2019 Microfabricated blood vessels for modeling the vascular transport barrier vol 14 (Springer US)

[25] Kirby B 2010 Micro-And Nanoscale Fluid Mechanics: Transport in Microfluidic Devices

[26] Wang Y I and Shuler M L 2018 UniChip enables long-term recirculating unidirectional perfusion with gravity-driven flow for microphysiological systems Lab Chip 18 2563–74

[27] Yang Y, Fathi P, Holland G, Pan D, Wang N S and Esch M B 2019 Pumpless microfluidic devices for generating healthy and diseased endothelia Lab Chip 19 3212–9

[28] Sonmez U M, Cheng Y W, Watkins S C, Roman B L and Davidson L A 2020 Endothelial cell polarization and orientation to flow in a novel microfluidic multimodal shear stress generator Lab Chip 20 4373–90

[29] Russo T A, Banuth A M M, Nader H B and Dreyfuss J L 2020 Altered shear stress on endothelial cells leads to remodeling of extracellular matrix and induction of angiogenesis PLoS One 15 1–17

[30] Noria S, Xu F, McCue S, Jones M, Gotlieb A I and Langille B L 2004 Assembly and Reorientation of Stress Fibers Drives Morphological Changes to Endothelial Cells Exposed to Shear Stress Am. J. Pathol. 164 1211–23

[31] Yang L, Shridhar S V, Gerwitz M and Soman P 2016 An in vitro vascular chip using 3D printing-enabled hydrogel casting *Biofabrication* 8 35015

[32] Tanaka T, Nakatani K, Morioka K, Urakawa H, Maruyama N, Kitagawa N, Katsuki A, Araki-Sasaki R, Hori Y, Gabazza E C, Yano Y, Wada H, Nobori T, Sumida Y and Adachi Y 2003 Nitric oxide stimulates glucose transport through insulin-independent GLUT4 translocation in 3T3-L1 adipocytes *Eur*. J. Endocrinol. 149 61–7

[33] Noris M, Morigi M, Donadelli R, Aiello S, Foppolo M, Todeschini M, Orisio S, Remuzzi G and Remuzzi A 1995 Nitric Oxide Synthesis by Cultured Endothelial Cells Is Modulated by Flow Conditions *Circ*. Res. 76 536–43

[34] Dubey R K, Gillespie D G, Mi Z and Jackson E K 2001 Endogenous cyclic AMP-adenosine pathway regulates cardiac fibroblast growth Hypertension 37 1095–100

[35] Godard M P, Johnson B A and Richmond S R 2005 Body composition and hormonal adaptations associated with forskolin consumption in overweight and obese men *Obes*. Res. 13 1335–43

[36] Gauthier M S, Miyoshi H, Souza S C, Cacicedo J M, Saha A K, Greenberg A S and Ruderman N B 2008 AMP-activated protein kinase is activated as a consequence of lipolysis in the adipocyte: Potential mechanism and physiological relevance *J*. Biol. Chem. 283 16514–24

[37] Perrot C Y, Sawada J and Komatsu M 2018 Prolonged activation of cAMP signaling leads to endothelial barrier disruption via transcriptional repression of RRAS FASEB J. 32 5793–812

[38] Mullins G R, Wang L, Raje V, Sherwood S G, Grande R C, Boroda S, Eaton J M, Blancquaert S, Roger P P, Leitinger N and Harris T E 2014 Catecholamine-induced lipolysis causes mTOR complex dissociation and inhibits glucose uptake in adipocytes *Proc*. Natl. Acad. Sci. 111 17450–5

[39] Niu W, Bilan P J, Hayashi M, Da Y and Yao Z 2007 Insulin sensitivity and inhibition by forskolin, dipyridamole and pentobarbital of glucose transport in three L6 muscle cell lines *Sci*. China Ser. C Life Sci. 50 739–47

[40] Loskill P, Sezhian T, Tharp K M, Lee-Montiel F T, Jeeawoody S, Reese W M, Zushin P J H, Stahl A and Healy K E 2017 WAT-on-a-chip: A physiologically relevant microfluidic system incorporating white adipose tissue Lab Chip 17 1645–54

[41] Rogal J, Binder C, Kromidas E, Roosz J, Probst C, Schneider S, Schenke-Layland K and Loskill P 2020 WAT-on-a-chip integrating human mature white adipocytes for mechanistic research and pharmaceutical applications *Sci*. Rep. 10 1–12

[42] Yang F, Carmona A, Stojkova K, Garcia Huitron E I, Goddi A, Bhushan A, Cohen R N and Brey E M 2021 A 3D human adipose tissue model within a microfluidic device Lab Chip 21 435–46

[43] Paek J, Park S E, Lu Q, Park K T, Cho M, Oh J M, Kwon K W, Yi Y S, Song J W, Edelstein H I, Ishibashi J, Yang W, Myerson J W, Kiseleva R Y, Aprelev P, Hood E D, Stambolian D, Seale P, Muzykantov V R and Huh D 2019 Microphysiological Engineering of Self-Assembled and Perfusable Microvascular Beds for the Production of Vascularized Three-Dimensional Human Microtissues ACS Nano 13 7627–43

[44] Li X, Xia J, Nicolescu C T, Massidda M W, Ryan T J and Tien J 2019 Engineering of microscale vascularized fat that responds to perfusion with lipoactive hormones Biofabrication 11

