## Supplementary information for "Microphysiological Modeling of Vascular Adipose Tissue for Multi-Throughput Applications"

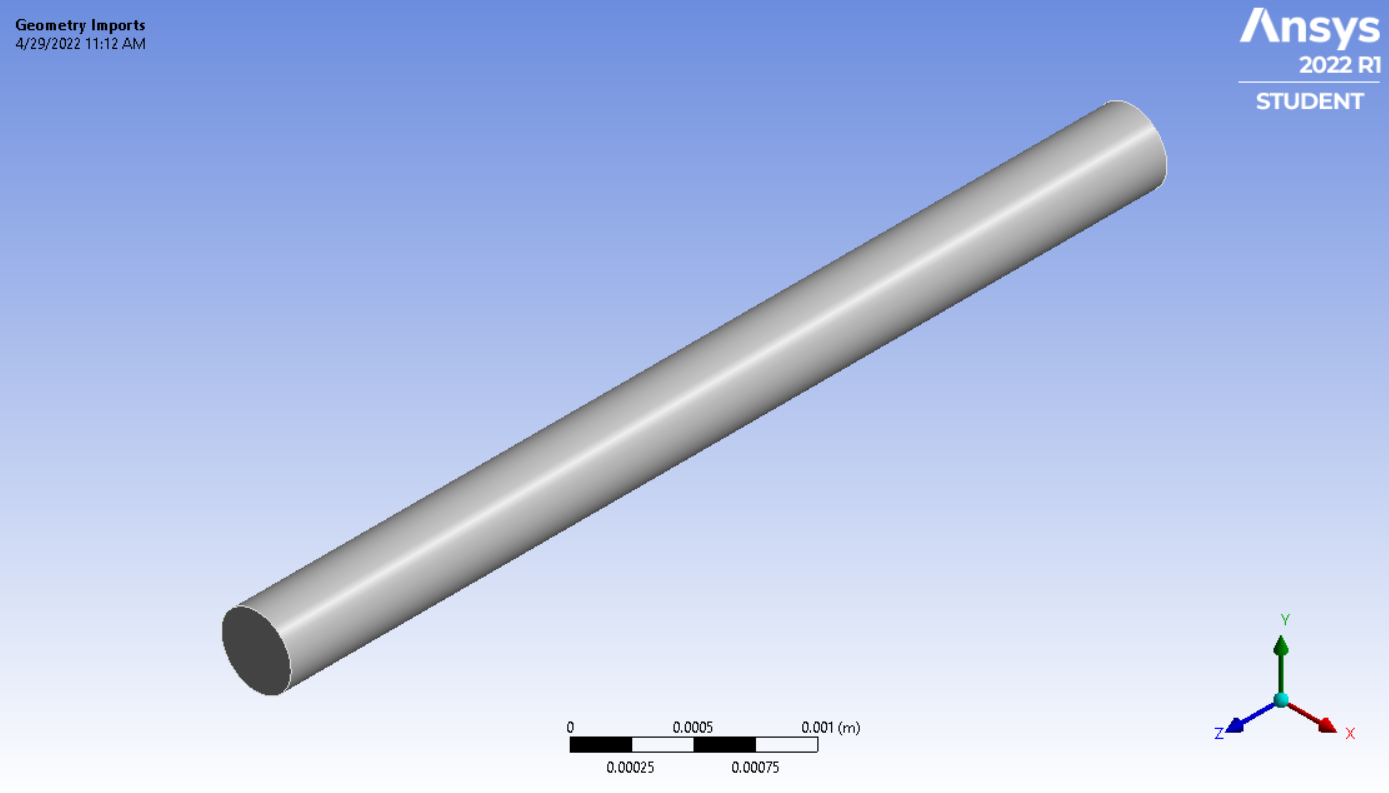

**Supplementary Figure S1. ANSYS design of vessel.** A 5 mm long cylinder with a diameter of 400 microns was created in ANSYS Design Modeler.

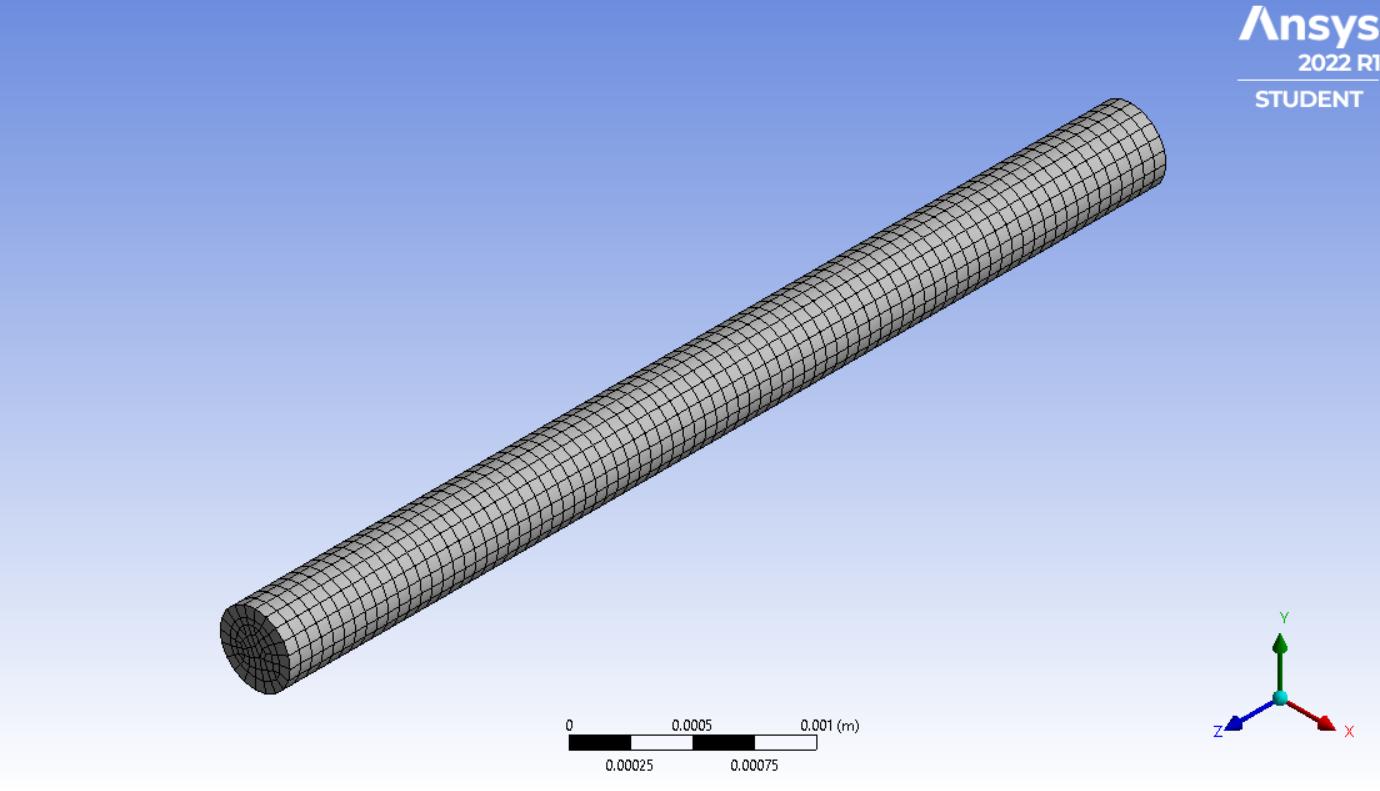

**Supplementary Figure S2. ANSYS meshing of vessel.** A linear mesh was created using a 0.1 mm mesh size.

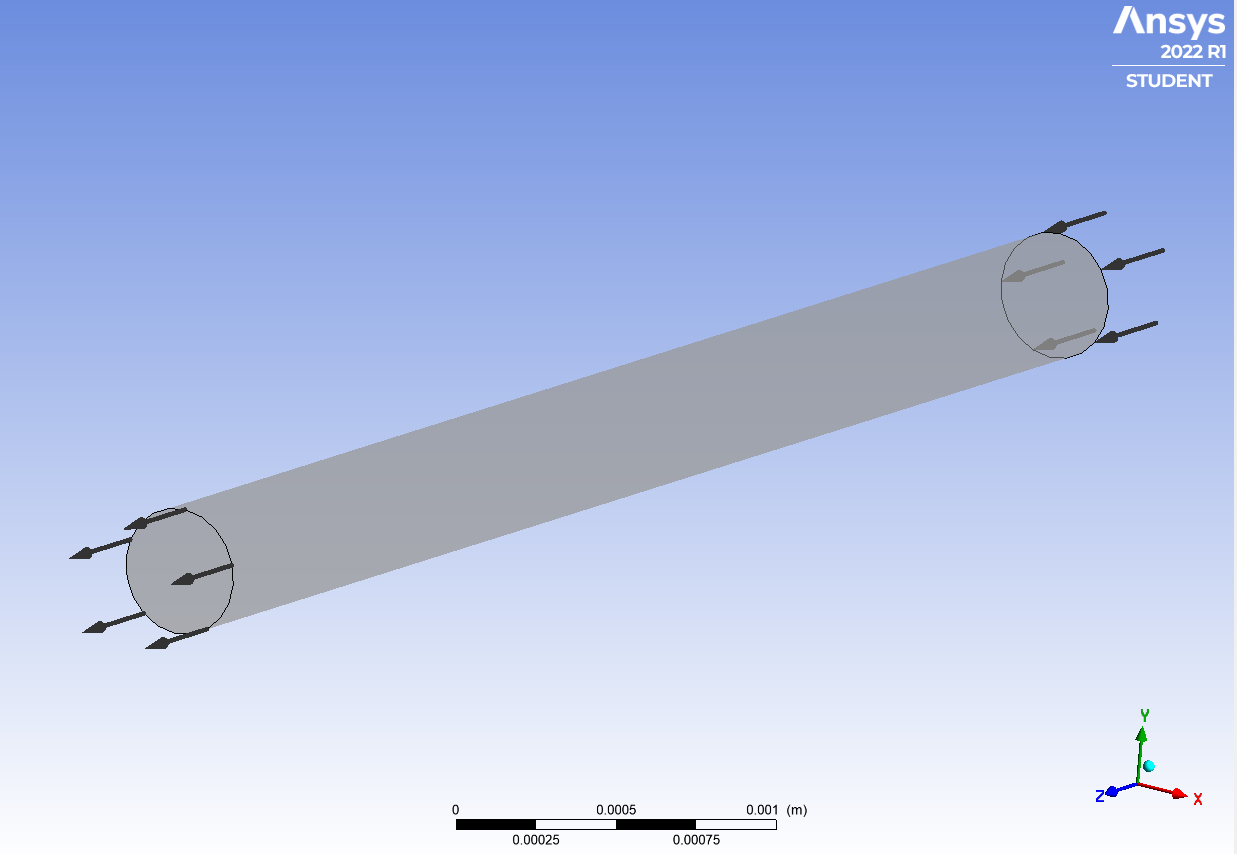

**Supplementary Figure S3. ANSYS boundary conditions of vessel.** The vessel was assigned an inlet, outlet, and wall.

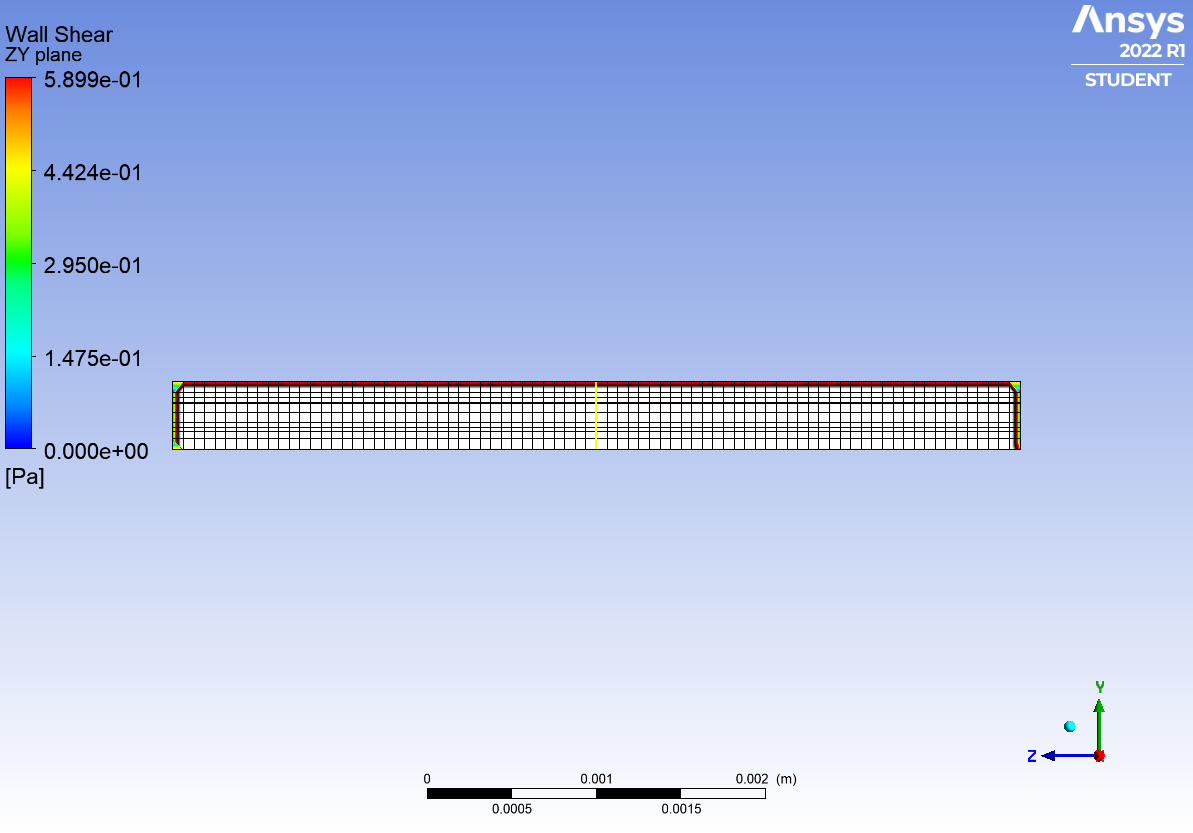

**Supplementary Figure S4. ANSYS CFD-Post of vessel.** In CFD-Post, after simulations have been completed, a 400-micron planar line was drawn in the y-direction across the diameter of the cylinder to plot the fluid velocity (V) and wall shear stress ($\tau)$.

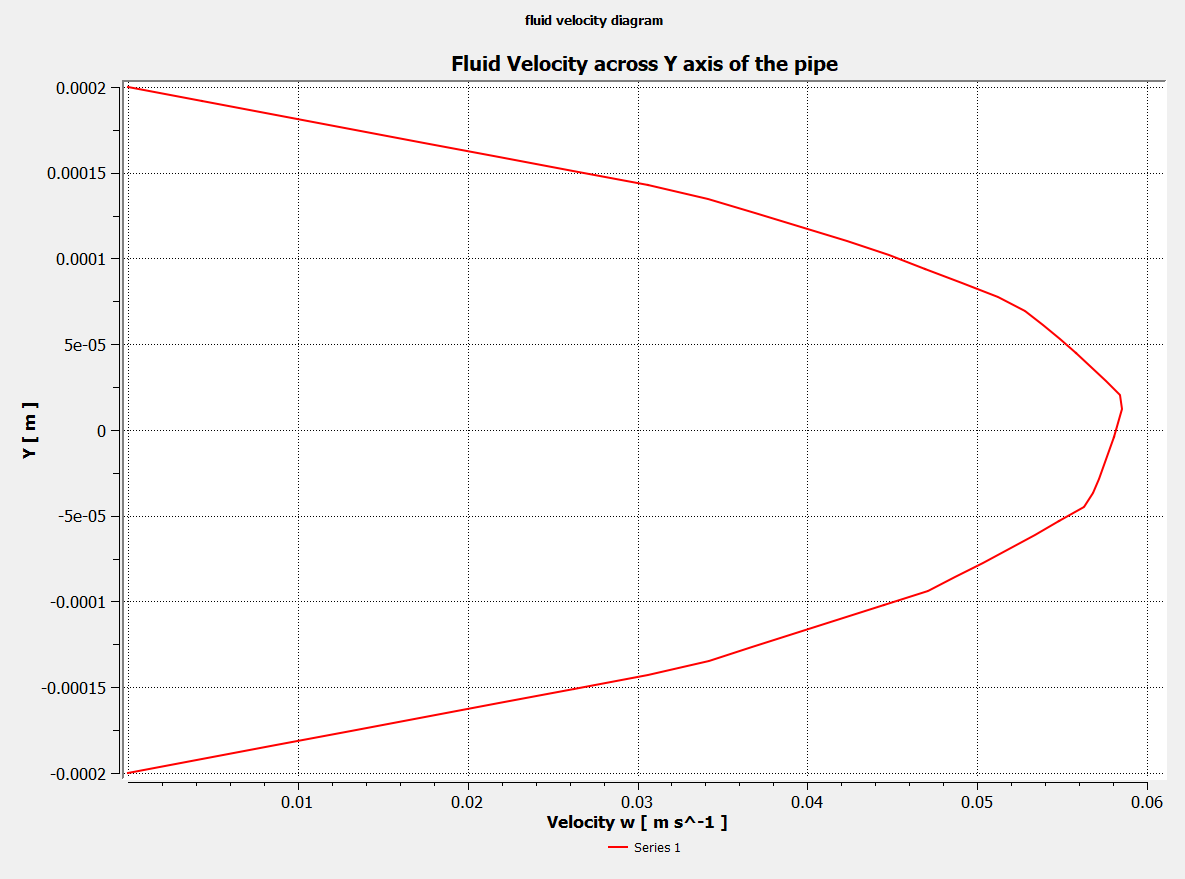

**Supplementary Figure S5. ANSYS CFD-Post: Fluid velocity within the vessel.** Fluid Velocity plotted as a function of distance. On the y-axis, 0 meters represents the center of the vessel. +/- 0.0002 meters represents of walls of the vessel.

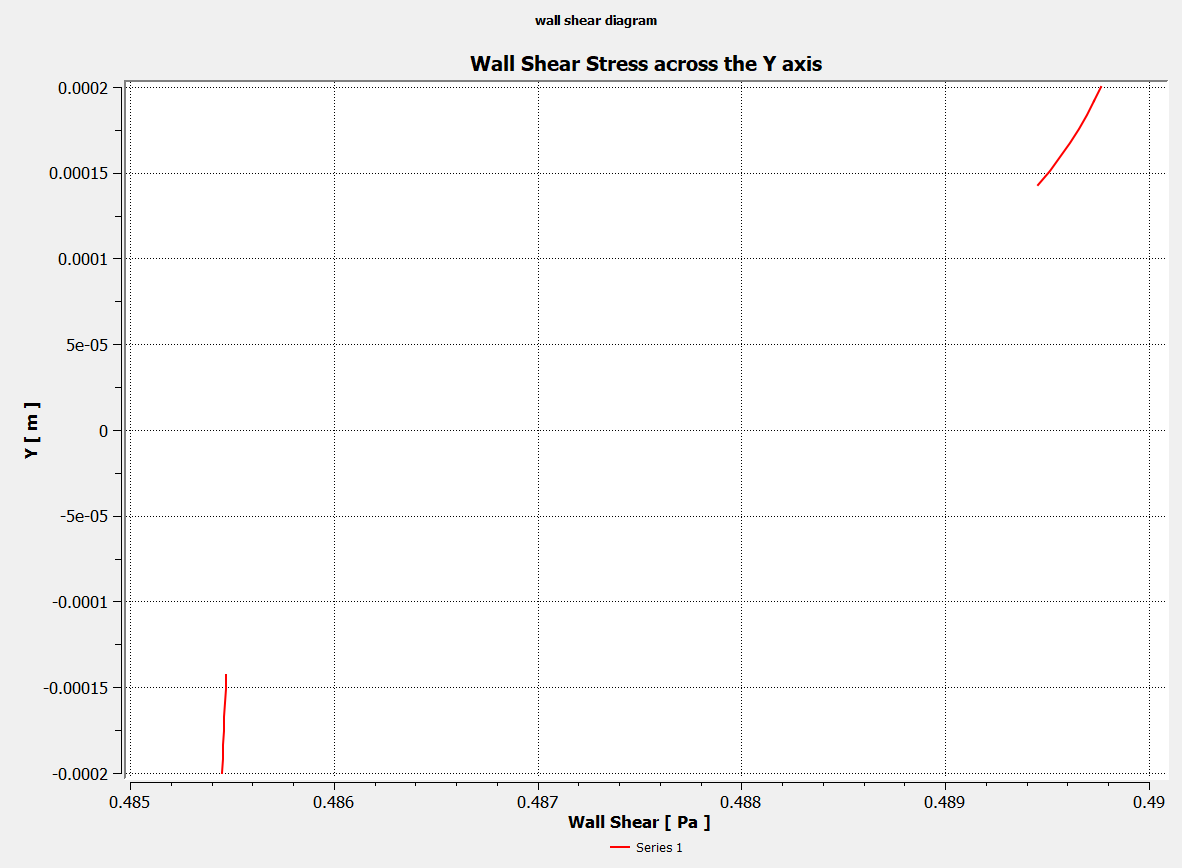

**Supplementary Figure S5. ANSYS CFD-Post: Wall shear stress within the vessel.** Wall shear stress plotted as a function of distance. On the y-axis, 0 meters represents the center of the vessel. +/- 0.0002 meters represents of walls of the vessel.

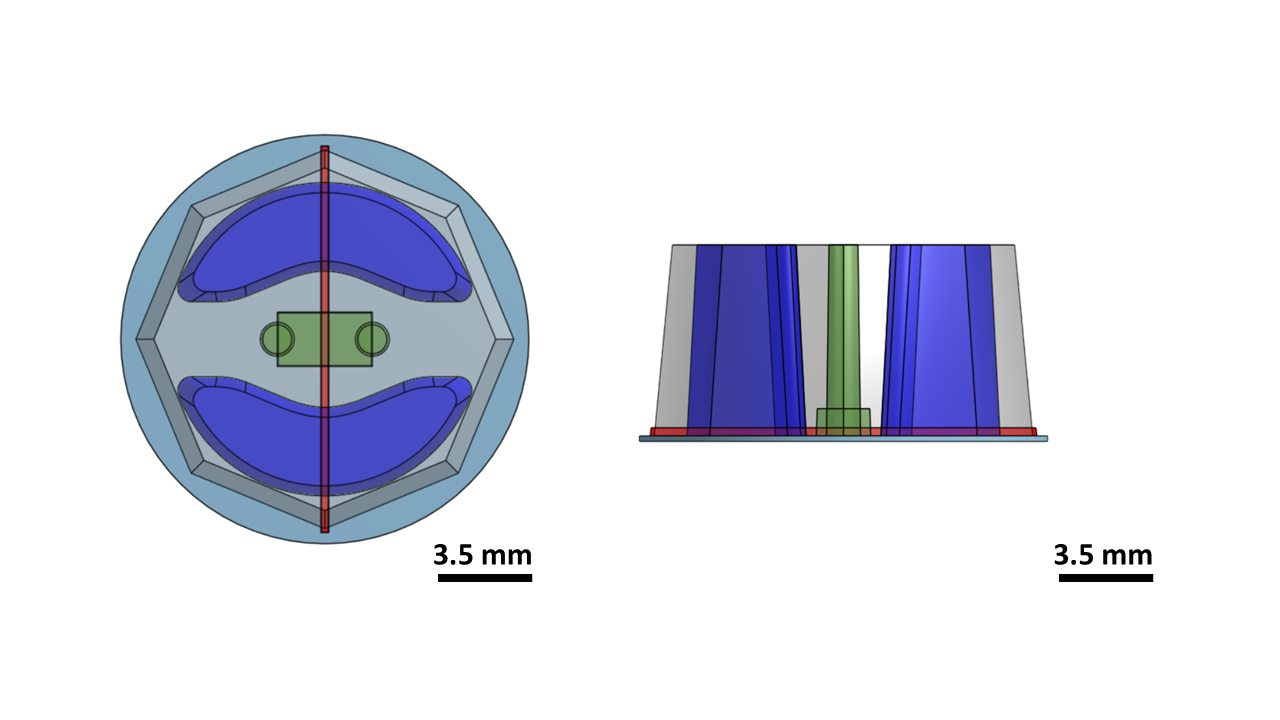
 **Supplementary Figure S7.** **Representative CAD design of the µAT model.**
